# Physical Linkage and Mate Preference Generate Linkage Disequilibrium for Behavioral Isolation in two Parapatric Crickets

**DOI:** 10.1101/468538

**Authors:** Thomas Blankers, Emma L. Berdan, R. Matthias Hennig, Frieder Mayer

## Abstract

Behavioral isolation is a potent barrier to gene flow and a source of striking diversity in the animal kingdom. However, it remains unclear if the linkage disequilibrium (LD) between sex-specific traits required for behavioral isolation results mostly from physical linkage between signal and preference loci or from directional mate preferences. Here, we test this in the field crickets *Gryllus rubens* and *G. texensis*. These closely related species diverged with gene flow and have strongly diverged songs and preference functions for the mate calling song rhythm. We map quantitative trait loci for signal and preference traits (pQTL) as well as for gene expression associated with these traits (eQTL). We find strong, positive genetic covariance between song traits and between song and preference. Our results show that this is in part explained by incomplete physical linkage: although both linked pQTL and eQTL couple male and female traits, major effect loci for different traits were never on the same chromosome. We suggest that the finely-tuned, highly divergent preference functions are likely an additional source of LD between male and female traits in this system. Furthermore, pleiotropy of gene expression presents an underappreciated mechanism to link sexually dimorphic phenotypes.

## INTRODUCTION

Behavioral isolation is often one the first and most potent forms of reproductive isolation to arise (Mayr 1963; Coyne and Orr 2004). This is somewhat paradoxical given that gene flow is often ongoing early on in speciation (Kirkpatrick and Ravigne 2002; Bolnick and Fitzpatrick 2007; Nosil 2008) and behavioral isolation typically requires linkage disequilibrium (LD) between at least two loci to be maintained. Gene flow and subsequent recombination threaten to break down this LD eroding isolation (Pinho and Hey 2010). Accordingly, the genetic architecture of behavioral isolation is a key feature that may predict the likelihood of speciation.

Often, LD needs to be maintained among multiple loci: Females may select males (or vice versa) based on multiple traits, each with different genetic underpinnings (Candolin 2003; Bro-Jorgensen 2010) and LD between these traits will increase divergence. Furthermore, both signal and preference are often polygenic (Bakker and Pomiankowski 1995; Ritchie and Phillips 1998; Gleason et al. 2002; Chenoweth and Blows 2006; Chenoweth and McGuigan 2010). The magnitude of LD required to maintain behavioral isolation in the face of gene flow is directly related to (1) the number of loci underlying signaling phenotypes (as well as the number of signaling phenotypes) and preferences, (2) their physical location in the genome, and (3) their effect sizes (Via and Hawthorne 1998; Coyne and Orr 2004; Arbuthnott 2009).

The theoretical literature has shown that speciation proceeds more readily if signal and preference loci are physically linked either via a single locus with pleiotropic effects or through close genomic proximity of separate signal and preference loci (Kirkpatrick and Hall 2004; Kopp et al. 2018). However, we have limited empirical insights into the genetic architecture of behavioral isolation. Current data provides evidence for linked signal and preference loci (pleiotropy or close linkage) in certain systems such as for color morphs and preferences in *Heliconis* butterflies (Kronforst et al. 2006; Merrill et al. 2011, 2018) and *Medaka* fish (Fukamachi et al. 2009), acoustic communication in *Laupala* crickets (Shaw and Lesnick 2009), pheromone signals and discrimination in *Drosophila melanogaster* (Marcillac et al. 2005), and morph and morph preference in *Erythrura* finches (Pryke 2010). However, other systems show unlinked signals and preference genes such as other aspects of *Drosophila melanogaster* mating success and mating preferences (Ting et al. 2001) and pheromone communication in moths (Smadja and Butlin 2009)

This natural variation in genetic architecture may be due in part to the shape of the preference function, which can strongly impact whether sexual selection aids or hinders divergence and thus the likelihood of speciation (Servedio and Boughman 2017; Kopp et al. 2018). Unimodal preference functions centered around the same mean value across diverging populations (Fig 1A) can lead to stabilizing sexual selection which impedes divergence (van Doorn et al. 2004; Weissing et al. 2011; Kopp et al. 2018) or even lead to homogenization of preference loci across populations (Servedio and Burger 2014). Strong physical linkage of these preference alleles with signal traits would mitigate these counterproductive effects during divergence and increase the likelihood of speciation (Servedio and Burger 2014). However, with open-ended (Fig 1B), relative preferences, or with strongly divergent unimodal preferences (Fig 1C), sexual selection can facilitate divergence and speciation will proceed more readily, especially when signaling traits are also ecologically relevant (Kondrashov and Kondrashov 1999; Doebeli 2005). Under such a model, strong physical linkage between preference and signal loci may not be necessary for divergence (Lande 1982). More empirical examples where we have knowledge of both the shape of preference functions as well as information about the genetic architecture of signals and preferences is needed to empirically test these theoretical results.

**Figure 1.**
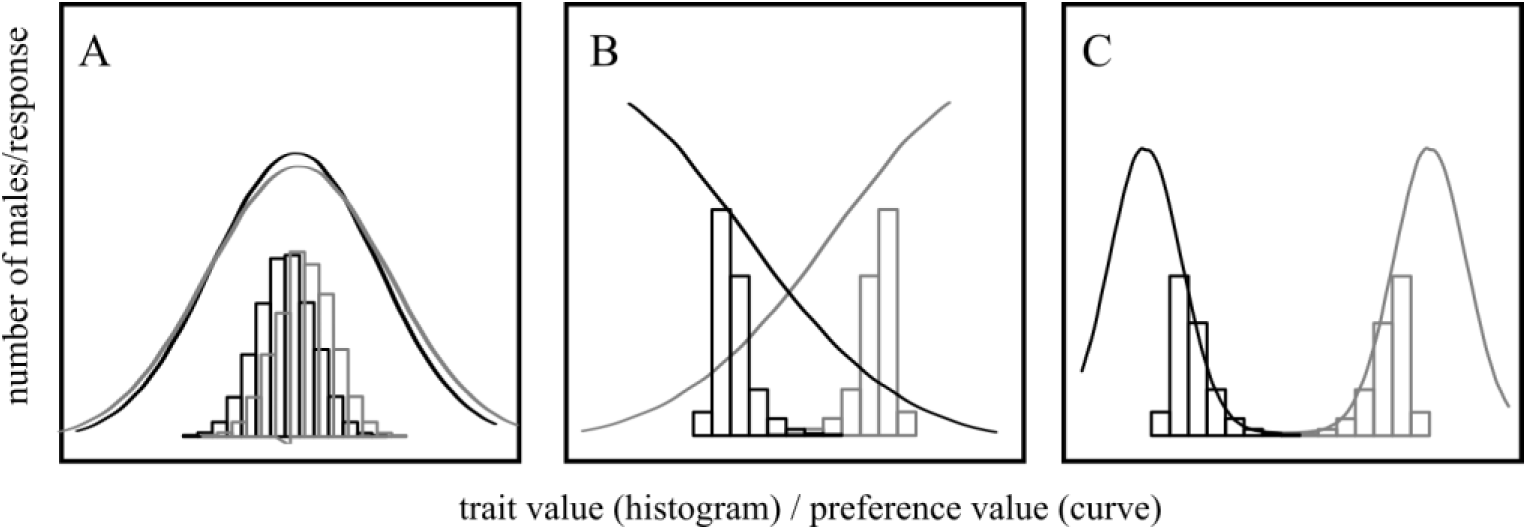
Schematic of male trait distribution and female preference function. Unimodal, non-divergent preferences lead to stabilizing selection (A). Open-ended (B) or strongly divergent (C) preference functions exert strong directional selection on the male trait, thereby creating genetic covariance even if loci reside on different (parts of) chromosomes.

Additional sources of variation in the genetic architecture may result from the fact that linkage will be detected more readily in systems where loci of relatively large phenotypic effect are linked among suites of traits or among signals and preferences. When more subtle aspects of the genetic architecture of co-evolving traits are linked, important associations might be missed using standard quantitative trait locus (QTL) methods. Behavioral variation between closely related species in general, and variation in sexually dimorphic traits contributing to behavioral barriers specifically, is strongly influenced by regulatory variation (Williams and Carroll 2009; Etges 2014; Mack and Nachman 2017) but the genetic architecture of expression and its role in speciation are widely underappreciated empirically and theoretically (Mack and Nachman 2017). One intriguing but unexplored example is that gene expression variation, due to the ubiquitous pleiotropy of regulatory variants (Chesler et al. 2005; Gibson and Weir 2005), could be a powerful means of generating additional LD between signal and preference if these traits have shared regulatory pathways or otherwise co-expressed loci. Although traditional QTL analysis may uncover these variants, integrating *p*QTL (i.e. with a behavioral, morphological, or physiological trait as the response variable) and *e*QTL (i.e. with the expression level of a gene or transcript as the response variable) is a powerful approach that may help to uncover additional loci of small effect if eQTL harbor mutations with weak effect on the phenotype but with sufficiently strong effects on gene expression related to that phenotype.

Here, we use QTL mapping to identify the number, distribution and effect size of loci associated with variation in the multivariate acoustic mate signal and with a major dimension of sexual selection resulting from female preference for the song rhythm in the field crickets *Gryllus rubens* and *G. texensis*. These sibling species are widely distributed across the eastern and southern USA (Alexander 1962; Walker 2017). Acoustic mate choice is a major driver of reproductive isolation (Walker 1998; Gray and Cade 2000; Gray 2005; Blankers et al. 2015a) which evidence suggests is strong: no natural hybrids have been documented and no females inseminated with heterospecific sperm have been collected (Gray and Cade 2000). Demographic analyses show that gene flow ceased roughly 18,000 years ago after initial divergence commenced 0.5 million years ago (Blankers et al. 2018b). Two male song traits that have diverged strongly between the species, pulse rate (i.e. the repetition rate of sound pulses) and carrier frequency (the pitch of the song), are both associated with unimodal preference functions. However, pulse rate preferences are finely tuned to the male song and strongly divergent among species, whereas carrier frequency preferences are broadly overlapping across species (Blankers et al. 2015b,a).

We hybridized wild-caught parental lines in the lab to obtain segregating mapping populations and looked for associations between transcriptome-wide SNP markers and variation in pulse rate and carrier frequency (pQTL scan). We then correlate phenotypic variation in the mapping population with variation in gene expression across more than 27,312 transcripts and perform an eQTL scan for all trait-associated transcripts. Our results on patterns of linkage among pQTL and eQTL significantly advance our understanding of the genetic architecture of behavioral isolation and provide important new insights into mechanisms of trait-preference co-evolution and divergence.

## MATERIAL & METHODS

Crickets were collected from allopatric locations [*G. texensis*: 84 females from Austin (TX), Lancaster (TX), and Round Rock (TX); *G. rubens*: 76 females from Gainesville (FL), Lake City (FL), and Live Oak (FL)] but patterns of reproductive isolation are similar across zones of sympatry and allopatry indicating that reinforcement is absent in this system (Izzo and Gray 2004). We generated eight mapping families encompassing all four possible types of backcrosses to pure *G. rubens* using parental individuals selected to maximize the potential phenotypic space for the hybrid offspring. Selected pairs were kept in the breeding boxes with water and food *ad libitum* and oviposition substrate for 1 week after the day the first eggs were recorded, after which both individuals were sacrificed and processed for RNA sequencing.

Measurements of the male song envelope (pulse rate) and spectrum (carrier frequency) were done using custom software (LabVIEW 2009) and the recording temperature (mean = 25.1 °C +/- 1.05 SD) was used to standardize the measurements. Female preferences were tested under dark and anechoic conditions, using a trackball system (Hennig et al. 2016): a Styrofoam sphere floating on pressurized air that can be easily moved by the cricket while infrared sensors underneath record the sphere’s movement in lateral and longitudinal directions. Custom software (LabVIEW, 2009) was used to present stimuli (Table S1) as well as negative (silence and pure frequency tones) and positive (highly attractive stimulus) controls and to analyze the feedback from the optical sensors. The lateral movement of a female during signal presentation was averaged between the consecutive playbacks from the two speakers (order of active and silent speaker was randomized across trials) and normalized with respect to the response to the attractive control signal.

Preferences were quantified in two ways: The stimulus with the highest phonotactic response was considered the peak preference score; The second approach quantified preference functions more broadly by projecting individual responses of all backcrosses to all eight stimuli onto a linear discriminant function (‘lda’ in the R-package ‘MASS’) (Venables and Ripley 2002), which had been trained on parental data (N = 73 *G. rubens* and N = 44 *G. texensis* females). This LD1 score will be referred to as pulse rate preference function from hereon and describes multiple aspects of female preference through the variable correlation of test patterns with the linear discriminant function (Table S2).

After phenotyping and/or crossing, each individual was played back its control stimulus for 10 minutes, preserved in RNAlater following the manufacturers recommendations, and transferred to −80°C. All libraries were sequenced on a HiSeq 2000 (Illumina, San Diego, California) at a depth of 13 libraries per lane with paired-end 100 bp reads. Reads were processed using Flexbar (Dodt et al. 2012) and transcript-level information was obtained by mapping the reads against the *G. rubens* reference transcriptome (Berdan et al. 2016) using Bowtie2 (Langmead and Salzberg 2012). SNPs were called using the Genome Analysis Toolkit (DePristo et al. 2011; Van der Auwera et al. 2013) and filtered using GATK and VCFtools (Danecek et al. 2011). Additional details on husbandry, phenotyping, and SNP calling and filtering are in the supplementary methods.

### Linkage mapping

We conducted a chi-square test for every SNP to determine if the segregation of alleles fit an autosomal or a sex-linked model (FDR corrected p < 0.1). We removed SNP loci if more than two families (out of eight) had missing genotype data and retained one SNP per transcript to avoid repetition. Exceptions were made for eight loci that were of special interest. Because crickets have XX-XO sex determination, only families with F1 dams have recombining sex chromosomes and only in a single family were we able to recover sufficient X-linked markers. All linkage and QTL mapping information for the X chromosome is thus based on that single family of 40.

Linkage maps were generated in Joinmap 4.1 (van Ooijen 2006) for each family individually. The total sample size was 288 (143 females and 145 males) and family sizes varied between 25 and 43. Linkage groups were created with a log-of-odds (LOD) threshold equal to 4.0 or 5.0. The Kosambi mapping function was used to convert recombination frequencies to centi-Morgans (cM). A consensus map was constructed using the map integration tool. Linkage groups from individual families were joined if they shared two or more markers.

### Heritability and genetic covariance

To estimate narrow-sense heritability of and genetic covariance among male signal traits and female preferences we used phenotypic data from grandparental and parental lines and their backcross offspring. We first fitted mixed models in lme4 (Bates et al. 2014) and estimated heritability using REML. We then fitted Bayesian Animal models in MCMCglmm (Hadfield 2010) using an inverse Wishart prior (Gelman 2006) and checked for autocorrelation, effective sample size, and chain convergence following the MCMCglmm course notes (Hadfield 2012). We then fitted multi-response models with male pulse rate, male carrier frequency, and female pulse rate preference as response variables and ran 1 million iterations discarding the first 100,000 as burn-in. The median and 95% Honest Posterior Density (HPD) interval of the heritability of each trait and genetic covariance (and correlation) between each trait pair were estimated from the posterior distribution.

We used similar models to estimate the heritability for each of the 27,312 transcripts, except for these models we only ran 100,000 iterations to accommodate computational resources. Due to the asymptotic patterns of some of the posterior distributions (approximating but not overlapping zero), we considered all transcripts with the lower tail of the 95% HPD interval higher than 0.01 to have non-zero heritability.

### pQTL mapping

The goal here was to establish the number and distribution of genetic loci contributing to variation in the main divergent phenotypes used in intersexual acoustic communication. We used R/QTL (Broman et al. 2003) in R (R Development Core Team 2016) to detect QTL for pulse rate, carrier frequency, and pulse rate peak preference and preference function (LD1 scores) at false discovery rate [FDR] < 5% (“significant”) or FDR < 63% (“suggestive”), following recommendations by the Complex Trait Consortium (Members of the Complex Trait Consortium 2003). We excluded two males for which song recordings did not meet minimal quality standards leaving 143 females and 142 males for pQTL mapping. We first used ‘scanone’ with Haley-Knott regression (Haley and Knott 1992) to identify the single strongest QTL for each trait, followed by 1,000 permutations to establish a significance threshold at the 5% and 63% level. We then used the multiple-QTL model approach (Broman and Sen 2009) to scan for additional QTL, refining QTL positions and establishing whether the model LOD score increased beyond the penalized LOD score threshold. The thresholds for FDR equal to 5% and 63% were obtained using 1,000 permutations of the ‘scantwo’ function. Cross type was included as a covariate in the models initially but removed if not significant. The magnitude of the additive effects and the 95% Bayesian credible interval was estimated from the model. To estimate the true number of loci underlying the phenotypic traits, we used a custom code based on (Otto and Jones 2000) to estimate the QTL detection threshold, the true number of loci, and the amount of missing variation given the results of our experiment. The code is available at github.com/thomasblankers/statistics/QTL_power_detect.r.

### eQTL mapping

The goals here were (i) to identify transcripts for which expression covaries with the main phenotypic traits used by males (pulse rate, carrier frequency) and females (pulse rate preference) in intersexual acoustic communication and (ii) to unravel the genetic architecture of the expression of these transcripts. Reads from all backcross individuals were separately aligned to the reference transcriptome using Bowtie (Langmead et al. 2009) and transcript abundances were calculated for each stage using RSEM (Li and Dewey 2011). We imported the read abundance data into R using ‘tximport’ (Soneson et al. 2015). We performed a differential expression analysis using a continuous model with both pulse rate and carrier frequency as fixed effects and cross as a covariate (expression ~ cross + pulse rate + carrier frequency) to account for cross effects and for the correlation between traits (see Results). We fit these models in DESeq2 (Love et al. 2014) with Wald’s test for significance. For pulse rate preference, we similarly fit a continuous DESeq model with cross as a covariate. For all models, we considered transcripts with a Benjamini-Hochberg (Benjamini and Hochberg 1995) corrected p-value < 0.01 to be significantly associated with the trait of interest.

We then performed a similar analysis using robust regression models in the R package limma (Ritchie et al. 2015). Here we used log2-TMM normalization to normalize the count data using the ‘calcNormFactors’ and ‘cpm’ function in edgeR. We then fitted linear models with robust regression and estimated empirical Bayes statistics for differential expression. All loci with adjusted P-value < 0.01 were considered significant.

For eQTL mapping, we kept only those loci that were significant in both the DESeq2 and the limma analysis and that had non-zero heritability. We retained two sets: one including all the above transcripts (“permissive” set) and the other containing those that have a relatively strong relationship with the trait (“conservative” set). The latter set consisted of transcripts that had partial η^2^ values (‘eta.square’ function in the R package heplots (Fox et al. 2018)) for their association with variation in the trait in the top 25% (for pulse rate η^2^ > 0.13, for carrier frequency η^2^ > 0.07, for pulse rate preference η^2^ > 0.13).

We used the ‘mqmscanall’ function on a trait-by-trait basis to perform a multiple eQTL scan for each transcript. We included 16 cofactors in the analysis, one for each linkage group at the median marker. We obtained LOD thresholds corresponding to FDR < 5% using 1,000 permutations of the “mqmscanall” function to establish significance of eQTL. For all significant eQTL, we checked if they were *cis* (regulatory substitution found on the same linkage group as the transcript itself) or *trans* (eQTL and corresponding transcript on different linkage groups) by comparing the eQTL location with the position of the transcript in the genome. To obtain linkage group level information about the location of as many transcripts as possible, we expanded our existing linkage map by including all loci that could be mapped to a linkage group (but not necessarily to a position within the group). This expanded map was only used to ascertain physical locations for eQTL transcripts.

To examine whether the transcripts that covaried in expression with male and female traits were also differentially expressed between species, we performed a differential expression analysis using the grandparents used to create the QTL mapping families as well as individuals previously sequenced (using similar methods as described above) for a population genetic study (Blankers et al. 2018b). Transcripts were considered differentially expressed if the adjusted p-value of Wald’s significance test was < 0.01 and read count differed at least 2-fold (log_2_-fold difference ≥ 1).

## RESULTS

### Phenotypes

Phenotypes were unimodally distributed within species or cross lines (Fig 2, Table 1). Values for first generation interspecific hybrids were intermediate but biased towards the maternal parent for pulse rate (*G. rubens* dam: 55.27 pulses s^−1^, *G. texensis* dam: 61.96 pulses s^−1^; t_22_ = −6.72; *P* < 0.0001) and carrier frequency (*G. rubens* dam: 4.82 kHz, *G. texensis* dam: 5.01 kHz; t_22_ = −3.9146; *P* = 0.0004). Backcross distributions were also unimodal and intermediate between interspecific hybrids and *G. rubens*. All traits follow expectations for polygenic, additive inheritance. In addition to the preference measurements used in the downstream analyses (i.e. the peak preference and the discriminant function score), the preference functions for pulse rate were unimodal in both species, F_1_ hybrids, and first-generation backcrosses (Fig S1).

**Figure 2.**
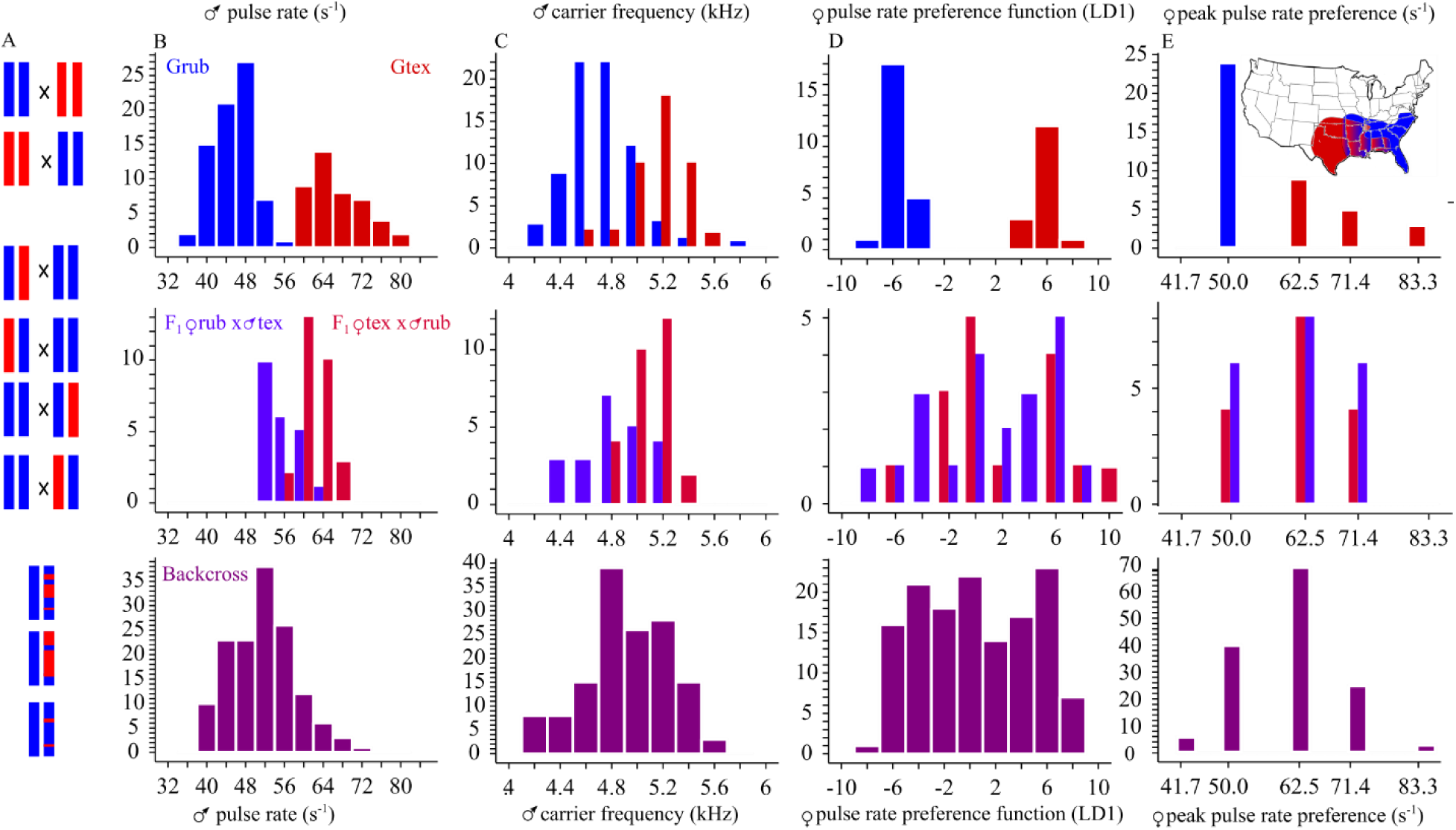
Phenotypic distributions. A: schematic crossing design. Diploid *G. rubens* (blue) and *G. texensis* (red) were crossed to obtain heterozygote first generation hybrid offspring in both cross directions. All possible combinations of hybrid-G*. rubens* were paired to create segregating backcross offspring. B-E: phenotypic distributions of parental (top panels), hybrid (middle panels), and backcross (bottom panels) offspring. Male pulse rate and carrier frequency are shown in B and C, female preference is shown in D (pulse rate preference, i.e. LD1 scores representing composite phonotactic scores on all 8 pulse rate test stimuli) and E (peak preference). The inset map in E shows the approximate geographic distribution of the parental species and their zone of overlap in the United States based on (Walker 2017).

**Table 1.**
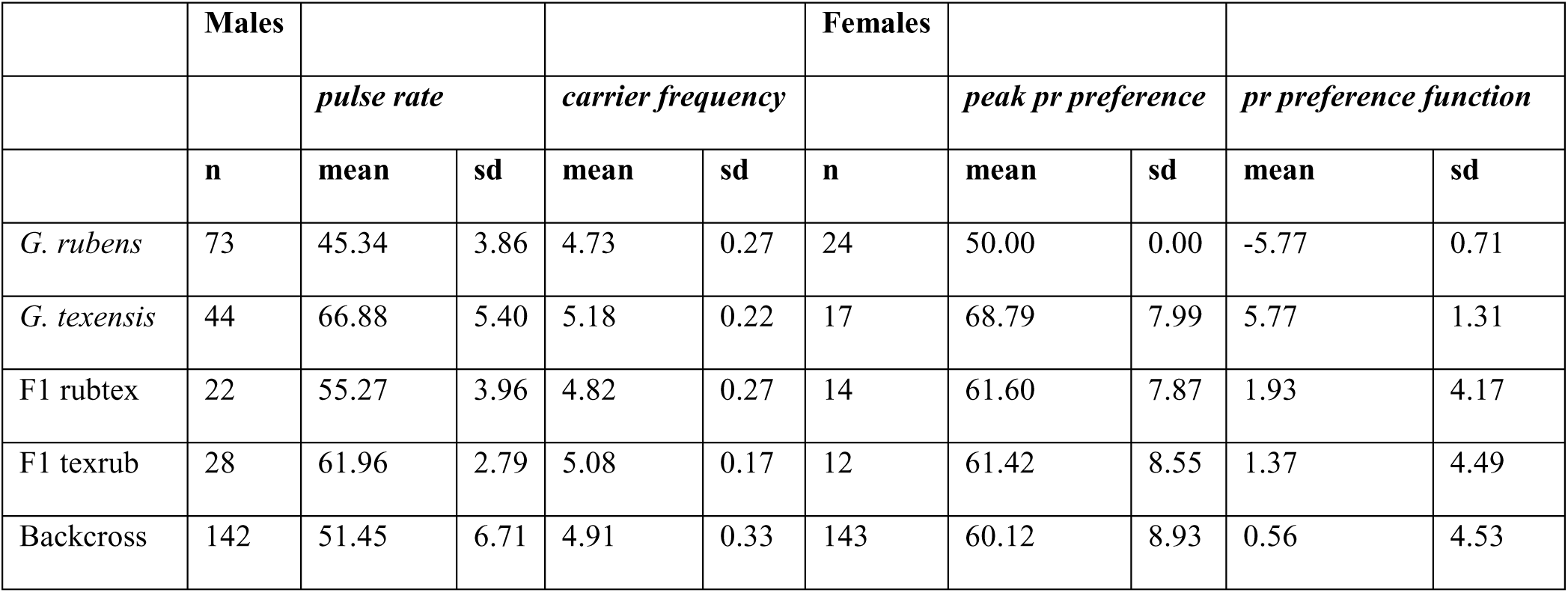
Phenotypic distributions of parental and hybrid generations. Pulse rate in pulses per second, carrier frequency in kilo Hertz, peak preference in pulses per second, and pulse rate preference function in dimensionless units of correlation with LD1. Rubtex are F_1_ indivduals with a *G. rubens* dam and texrub are F_1_ individuals with a *G. texensis* dam

### Linkage mapping

We placed a total of 330 markers on our genetic map (Table S3). The markers were grouped in 15 autosomal linkage groups (LG), one more than the number of autosomes for *G. rubens* (Yoshimura 2005), and an X-linked group with a total map distance of 254.4 cM, an average marker spacing of 0.81 cM, and a maximum marker spacing of 14.40 cM. Linkage groups varied in length from 0.99 cM to 41.5 cM (mean 17.3 cM).

### Heritability and genetic covariance

REML heritabilities estimated from sire variance were 0.91, 0.51, and 0.61 for pulse rate, carrier frequency, and pulse rate preference, respectively. The Bayesian Animal models gave similar results with 95% HPD between 0.48 and 0.99, 0.49 and 0.74, and 0.27 and 0.99, respectively. Correlations among the traits were high: median correlations were 0.49 for corr(pulse rate, carrier frequency), 0.92 for corr(pulse rate, pulse rate preference), and 0.46 for corr(carrier frequency, pulse rate preference); 95% HPD intervals did not overlap with zero (0.24 – 0.70; 0.58 – 0.99; 0.16 – 0.71). All genetic covariances were positive, indicating that an increase in one trait was associated with an increase in the other trait.

### pQTL mapping

Using single interval mapping, we detected only a single significant QTL for each trait, except for peak preference for which both an autosomal and an X-linked QTL were significant at α = 0.05 (Fig S2). Because single QTL scans have limited power to detect small effect QTL, we added the significant QTL identified in single interval mapping to a multiple QTL model (MQM) and proceeded to scan for additional QTL at 5% (i.e. significant QTL) and 63% (i.e. suggestive QTL) FDR. In the final MQM (Fig 3; Table 2) we identified one significant and 4 suggestive pQTL for pulse rate (all on autosomes), 2 significant autosomal and 1 suggestive X-linked pQTL for carrier frequency, 2 significant and 2 suggestive autosomal pQTL for the pulse rate preference function (LD1 scores), and 1 significant autosomal and 1 significant X-linked as well as 2 suggestive autosomal QTL for peak pulse rate preference.

**Figure 3.**
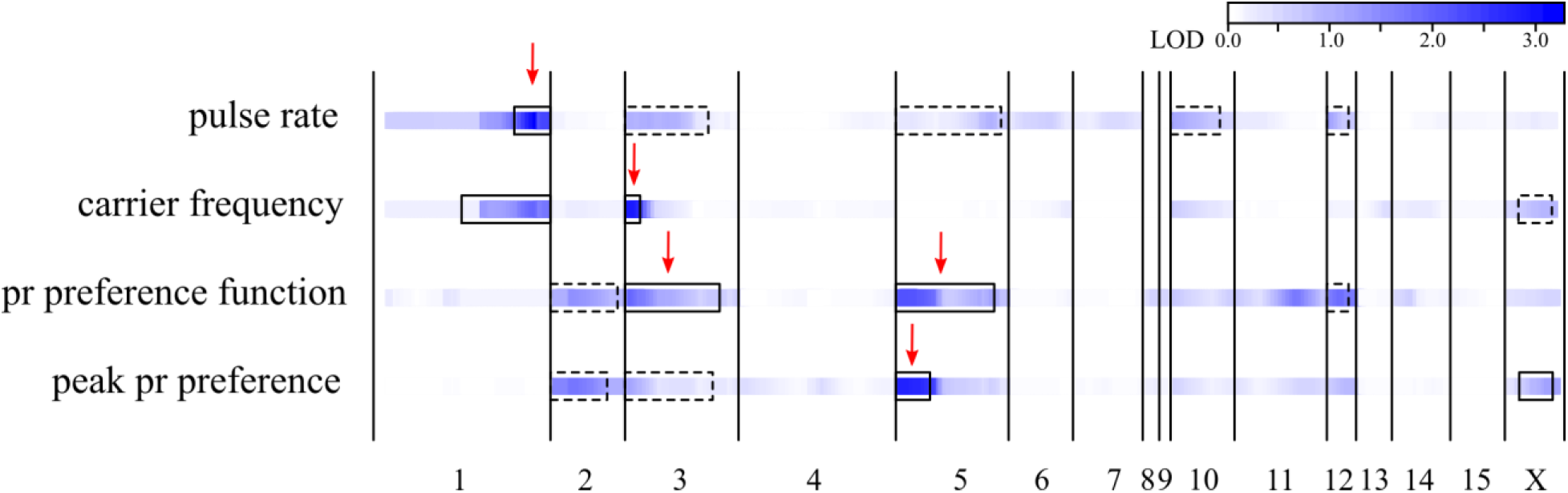
pQTL scan. For each of the four traits, the LOD scores along the 16 linkage groups is shown by the intensity of blue hues. The scale is shown on the top right. 95% Bayesian confidence intervals for significant (solid) and suggestive (dashed) are shown as boxes projected onto the heatmap. Red arrows indicate pQTL explaining > 10% of the backcross variance. “pr” abbreviates pulse rate. For single QTL interval mapping see Fig S2.

**Table 2.** pQTL effects. For each trait, the linkage group, the location, the nearest marker, the LOD score, the genotypic effects, and the pQTL effects expressed in trait mean change, number of standard deviations in *G. rubens* and percentage of backcross variance of each of the pQTL effects is shown. Significant pQTL (< 5% FDR based on penalized LOD score improvement of the multiple QTL model) are in bold. All pQTL effects are significantly larger than zero: \**P* < 0.05; † *P* < 0.01; ‡ *P* < 0.001

| Trait | LG | pQTL location (cM) | Nearest Marker | LOD | AA | AB | QTL effect (trait mean +/- SE) | % species difference | % backcross variance |
| --- | --- | --- | --- | --- | --- | --- | --- | --- | --- |
| **pulse rate** | **1** | 39.4 | c214087_g2_i1 | 3.21 | 48.72 | 51.83 | 3.62 +/- 0.94‡ | 17.24 | 14.8 |
|  | 3 | 6 | c215368_g2_i3 | 1.04 | 49.85 | 50.70 | 2.11 +/- 0.98* | 10.03 | 4.6 |
|  | 5 | 25 | c218669_g2_i3 | 0.97 | 49.14 | 51.32 | 2.15 +/- 1.05* | 10.23 | 4.3 |
|  | 10 | 0 | c186619_g1_i1 | 1.14 | 49.31 | 51.39 | 2.33 +/- 1.06* | 11.09 | 5.1 |
|  | 12 | 0 | c204487_g2_i1 | 1.22 | 49.11 | 51.23 | 2.18 +/- 0.94* | 10.37 | 5.5 |
|  | cross |  |  | 2.99 |  |  | 1.47 +/- 0.39‡ | 37.92 | 13.8 |
| **carr. freq.** | **1** | 39 | c214087_g2_i1 | 1.93 | 4.84 | 4.98 | 0.16 +/- 0.05† | 34.89 | 8.9 |
|  | **3** | 1 | c203593_g1_i1 | 2.89 | 4.83 | 5.00 | 0.19 +/- 0.05‡ | 41.76 | 13.4 |
|  | X | 9.9 | c205832_g2_i1 | 0.90 | 4.86 | 4.98 | 0.14 +/- 0.06* | 30.20 | 4.1 |
| **pulse rate pref. func. (LD1)** | **2** | 4.4 | c142606_g1_i1 | 1.63 | -0.38 | 1.32 | 1.84 +/- 0.68† | 15.94 | 7.4 |
|  | **3** | 0.1 | c218168_g2_i2 | 2.28 | -0.65 | 1.31 | 2.19 +/- 0.67† | 18.96 | 10.5 |
|  | **5** | 2 | c217193_g1_i1 | 2.35 | -0.49 | 1.37 | 2.20 +/- 0.67† | 19.07 | 10.8 |
|  | 12 | 2 | c203868_g1_i1 | 1.91 | -0.6 | 1.69 | 2.08 +/- 0.70† | 18.03 | 8.7 |
|  | cross |  |  | 2.7 |  |  | 0.53 +/- 0.15‡ | 4.57 | 12.4 |
| **peak pulse rate pref.** | **2** | 5.1 | c214277_g1_i1 | 1.94 | 58.29 | 61.56 | 3.99 +/- 1.35† | 21.23 | 8.8 |
|  | 3 | 0.1 | c218168_g2_i2 | 1.35 | 58.25 | 61.20 | 3.30 +/- 1.36* | 17.55 | 6.1 |
|  | **5** | 0 | c212100_g1_i1 | 3.00 | 57.52 | 62.17 | 5.07 +/- 1.36‡ | 26.96 | 13.9 |
|  | **X** | 9.8 | c205832_g2_i1 | 1.08 | 57.35 | 62.56 | 3.94 +/- 1.61* | 20.99 | 4.8 |
|  | cross |  |  | 2.17 |  |  | 0.94 +/- 0.30‡ | 5.02 | 9.9 |

The QTL for peak preference and preference function were largely overlapping, excepting an X-linked QTL for peak preference and a suggestive QTL on LG 12 for preference function (Fig 3, Fig S2). Carrier frequency and pulse rate also mapped to similar regions, with significant QTL overlapping on LG 1, and a suggestive pulse rate QTL on LG 3 overlapping with a significant QTL for carrier frequency. There was also QTL co-localization between male and female traits on LG 3 (all traits, QTL for pulse rate is suggestive), LG 5 (suggestive QTL for pulse rate, significant QTL for preference), LG 12 (suggestive QTL for both pulse rate and preference), and the X chromosome (carrier frequency and peak pulse rate preference (Fig 3, Fig S2).

The effect sizes for each QTL are shown in Table 2. For pulse rate, haploid allelic effects from 5 loci explained a total of 12.39 pulses per second, or 34.3% of the backcross variance. For carrier frequency this was 0.44 kHz or 26.4% of the variance across three loci and for peak preference and pulse rate preference function the total of four QTL effects was 16.07 pulses s^−1^ and 7.81 or 37.4% and 33.6% of the backcross variance, respectively. The combined effect size expressed as percentage of the difference between parental mean phenotypic values is much larger, but we note that these estimates are biased upwards due to our selective breeding of individuals from the extremes of the distributions. All QTL effects were significant (p-value for one sample t-test < 0.05) and of the same sign (i.e. *G. texensis* alleles always increase the trait values). Cross type effects were significant for all traits except carrier frequency but are not included in the sum of haploid allelic effects.

Using equation 6 in Otto & Jones (2000), we estimated the true number of loci [95% confidence interval] to be 23.30 [8.36-50.1] for pulse rate, 7.25 [2.61-15.59] for carrier frequency, 9.08 [2.83 – 21.11] for pulse rate peak preference, and 18.22 [5.66 – 42.32] for pulse rate preference function.

### eQTL mapping

We identified 430, 35, and 26 transcripts for which expression covaried with pulse rate, carrier frequency, and pulse rate preference (LD1), respectively (Table S4, S5, S6). The average narrow-sense heritability, *h*̅^2^, of these transcripts was 0.43 (*h*̅^2^ = 0.37 for all 27,312 transcripts) with a minimum of 0.00. After removing 69, 2, and 1 transcript(s) with very low heritability (lower 95% HPD interval < 0.01), *h*̅^2^ = 0.49 with a minimum of 0.07. The partial η^2^ for trait variation explained varied between 0.05 and 0.23 when considering all transcripts with non-zero heritability (“permissive” set) and between 0.12 and 0.23 when considering only the transcripts in the top 25% for the magnitude of trait association (“conservative” set; 109, 10, and 8 transcripts for pulse rate, carrier frequency, and pulse rate preference respectively). Some of these transcripts (43 out of 430 transcripts for pulse rate, 12 out of 35 for carrier frequency, and 5 out of 26 for preference) were also differentially expressed between the pure species (Table S4, S5, S6, Fig S3).

eQTL were significant between LOD > 3.0 and LOD > 2.0 depending on the trait and set of transcripts. We detected a total of 56 significant eQTL, 15 of which were from the conservative set of trait-associated transcripts, the remaining 41 from the permissive set. Of these, 39 from the permissive (6 from the conservative) transcripts covaried with pulse rate, 6 (4) with carrier frequency, and 11 (5) with pulse rate preference (Fig 4, Table 3).

**Figure 4.**
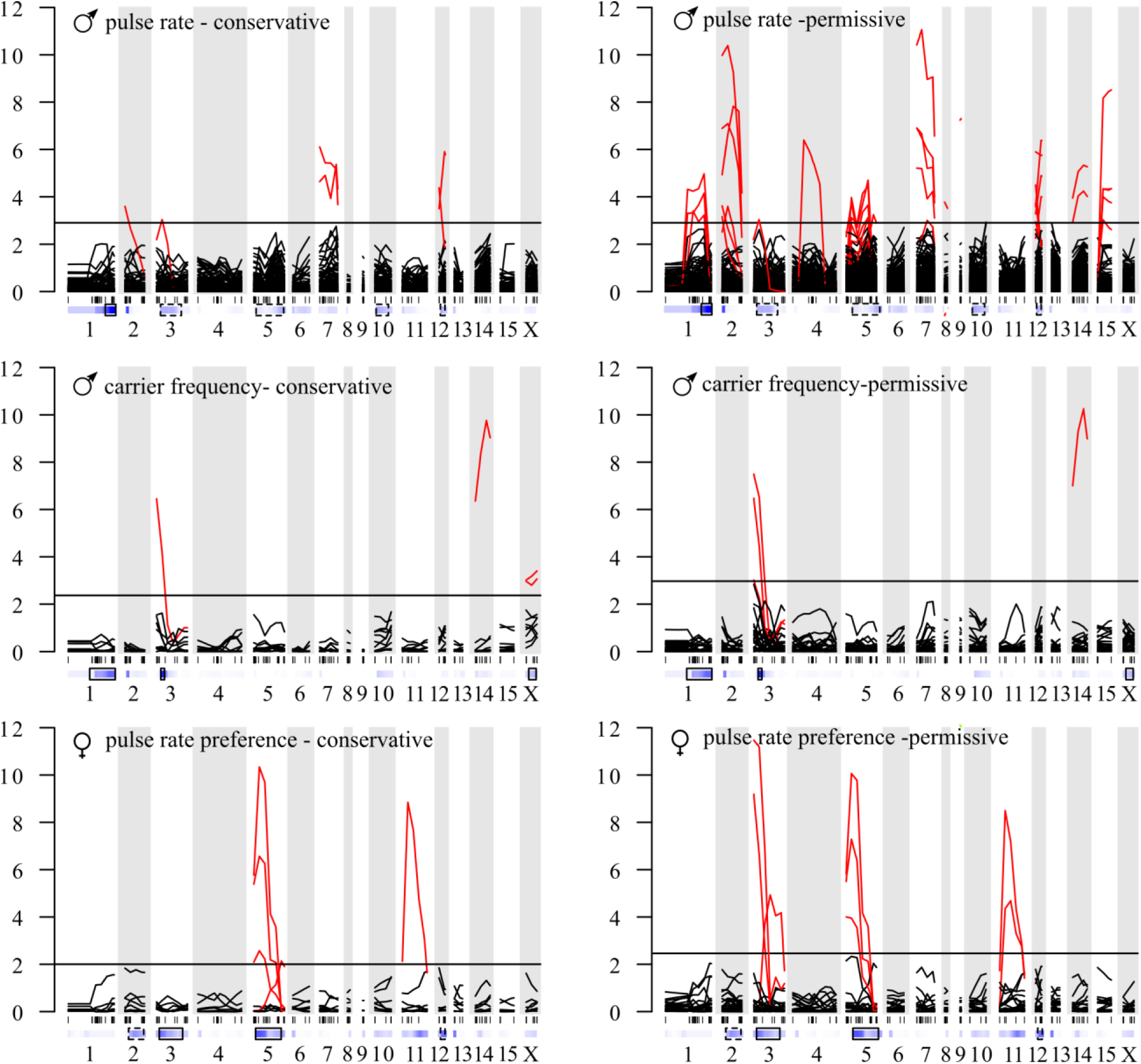
eQTL scan. The LOD score traces from the multiple QTL-model (one co-factor per linkage group) are shown for all transcripts that significantly covaried with pulse rate (top panels), carrier frequency (middle panels), and pulse rate preference (bottom panels). The horizontal black solid line shows the LOD threshold above which the false discovery rate is below 5% (see table 3). All significant eQTL are shown in red. The heatmap insets below each panel show the pQTL results (Fig 3) for comparison. See also Fig S4-S6 for transcript specific eQTL scan results.

**Table 3.** eQTL locations. For each transcript with a significant eQTL, the linkage group, position, and strength of association (LOD) are shown. The next three columns show the trait with which the expression level of the transcript covaries, the strength of the correlation between trait and transcript expression (partial η^2^ after accounting for covariates), and, if known, whether the transcript and the corresponding eQTL are located on the same linkage group (*cis* eQTL) or on different linkage groups (*trans* eQTL).

| Transcript | LG | eQTL location | LOD | Trait | Eta.sq | Set | Cis/trans |
| --- | --- | --- | --- | --- | --- | --- | --- |
| c220124_g1_i1 | 1 | 25.0 | 4.14 | pr | 0.09 | permissive | unkown |
| c219632_g1_i1 | 1 | 35.0 | 4.01 | pr | 0.12 | permissive | unkown |
| c205485_g2_i1 | 1 | 40.0 | 6.06 | pr | 0.09 | permissive | unkown |
| **c210636_g2_i5** | **2** | **0.0** | **3.59** | **pr** | **0.18** | **conservative** | **unkown** |
| c220300_g2_i7 | 2 | 0.0 | 3.54 | pr | 0.06 | permissive | unkown |
| c205742_g2_i4 | 2 | 5.0 | 3.72 | pr | 0.09 | permissive | unkown |
| c210780_g1_i1 | 2 | 5.0 | 4.03 | pr | 0.11 | permissive | unkown |
| c220753_g1_i3 | 2 | 5.0 | 10.92 | pr | 0.07 | permissive | unkown |
| c221848_g2_i3 | 2 | 5.0 | 7.50 | pr | 0.09 | permissive | cis |
| c210131_g1_i2 | 2 | 17.8 | 9.50 | pr | 0.13 | permissive | unkown |
| c205841_g4_i1 | 3 | 0.0 | 7.49 | cf | 0.10 | permissive | unkown |
| c207012_g2_i2 | 3 | 0.0 | 11.47 | pr pref | 0.13 | permissive | cis |
| c208039_g1_i2 | 3 | 0.0 | 3.02 | cf | 0.09 | permissive | unkown |
| **c209571_g2_i2** | **3** | **0.0** | **6.45** | **cf** | **0.14** | **conservative** | **unkown** |
| c218948_g2_i2 | 3 | 0.0 | 9.19 | pr pref | 0.13 | permissive | unkown |
| **c214456_g6_i2** | **3** | **5.0** | **3.04** | **pr** | **0.18** | **conservative** | **cis** |
| c206394_g1_i1 | 3 | 15.0 | 4.93 | pr pref | 0.14 | permissive | unkown |
| c220970_g1_i3 | 4 | 10.0 | 6.15 | pr | 0.12 | permissive | unkown |
| c199673_g1_i1 | 5 | 0.0 | 4.27 | pr | 0.12 | permissive | unkown |
| c216986_g2_i1 | 5 | 0.0 | 4.39 | pr | 0.11 | permissive | unkown |
| c220530_g1_i1 | 5 | 0.0 | 4.00 | pr pref | 0.12 | permissive | unkown |
| **c203543_g1_i1** | **5** | **5.0** | **2.57** | **pr pref** | **0.15** | **conservative** | **unkown** |
| **c208172_g4_i1** | **5** | **5.0** | **6.57** | **pr pref** | **0.14** | **conservative** | **unkown** |
| **c219551_g1_i2** | **5** | **5.0** | **10.34** | **pr pref** | **0.15** | **conservative** | **unkown** |
| c202635_g10_i | 5 | 20.0 | 4.54 | pr | 0.10 | permissive | unkown |
| c202662_g4_i1 | 5 | 20.0 | 3.80 | pr | 0.09 | permissive | unkown |
| c213768_g1_i2 | 5 | 20.0 | 3.32 | pr | 0.08 | permissive | unkown |
| c216251_g3_i4 | 5 | 20.0 | 4.42 | pr | 0.09 | permissive | unkown |
| **c218893_g1_i1** | **5** | **25.0** | **2.14** | **pr pref** | **0.16** | **conservative** | **unkown** |
| **c220971_g2_i1** | **7** | **0.0** | **6.11** | **pr** | **0.13** | **conservative** | **unkown** |
| c210495_g1_i1 | 7 | 5.0 | 8.97 | pr | 0.12 | permissive | cis |
| c218100_g2_i1 | 7 | 5.0 | 5.82 | pr | 0.10 | permissive | unkown |
| c201543_g3_i7 | 7 | 15.0 | 3.08 | pr | 0.09 | permissive | unkown |
| c205462_g3_i6 | 7 | 15.0 | 3.12 | pr | 0.13 | permissive | unkown |
| **c211026_g6_i1** | **7** | **15.0** | **5.37** | **pr** | **0.17** | **conservative** | **unkown** |
| c220188_g1_i1 | 8 | 2.4 | 8.03 | pr | 0.12 | permissive | cis |
| c205911_g1_i3 | 9 | 0.0 | 5.69 | pr | 0.07 | permissive | unkown |
| c208857_g7_i1 | 9 | 0.0 | 12.11 | pr pref | 0.12 | permissive | unkown |
| **c221227_g1_i2** | **11** | **5.0** | **8.84** | **pr pref** | **0.16** | **conservative** | **unkown** |
| c210974_g1_i1 | 11 | 10.0 | 4.68 | pr pref | 0.14 | permissive | unkown |
| c213294_g1_i2 | 12 | 0.0 | 3.21 | pr | 0.12 | permissive | unkown |
| **c213895_g1_i1** | **12** | **0.0** | **4.38** | **pr** | **0.18** | **conservative** | **unkown** |
| c220585_g1_i1 | 12 | 0.0 | 4.65 | pr | 0.09 | permissive | unkown |
| **c204736_g4_i1** | **12** | **5.0** | **5.91** | **pr** | **0.14** | **conservative** | **unkown** |
| c213099_g1_i1 | 12 | 5.0 | 3.67 | pr | 0.12 | permissive | unkown |
| c214279_g2_i1 | 12 | 5.0 | 4.13 | pr | 0.10 | permissive | unkown |
| c214910_g1_i1 | 14 | 10.0 | 5.64 | pr | 0.10 | permissive | unkown |
| c221673_g2_i1 | 14 | 10.0 | 4.28 | pr | 0.09 | permissive | unkown |
| **c222074_g1_i1** | **14** | **10.0** | **9.76** | **cf** | **0.19** | **conservative** | **cis** |
| c220034_g1_i1 | 14 | 13.4 | 3.07 | pr | 0.15 | permissive | unkown |
| c199619_g1_i1 | 15 | 0.0 | 3.58 | pr | 0.12 | permissive | unkown |
| c219073_g3_i4 | 15 | 5.0 | 3.64 | pr | 0.12 | permissive | unkown |
| c201025_g1_i1 | 15 | 10.0 | 3.84 | pr | 0.12 | permissive | cis |
| c214895_g3_i1 | 15 | 10.0 | 6.82 | pr | 0.08 | permissive | unkown |
| **c206857_g1_i1** | **X** | **9.9** | **3.06** | **cf** | **0.16** | **conservative** | **unkown** |
| **c213596_g1_i1** | **X** | **9.9** | **3.40** | **cf** | **0.17** | **conservative** | **unkown** |

Expanding the linkage map used for QTL mapping to include any transcript of known linkage group (but potentially unknown position within linkage groups due to limited shared markers among families) resulted in 1,611 transcripts with linkage group assignment (Table S7). However, the majority of the transcripts for which an eQTL was identified did not have a linkage group assigned and therefore information about *cis* versus *trans* regulation is limited; however, all 7 eQTL for transcripts of known location were *cis*-regulated (Table 3).

We find significant eQTL on most linkage groups, except LG 6, 10, and 13 (Fig 4, Table 3). There is a trend for eQTL to co-localize as 40/56 eQTL co-localized with at least one other eQTL (Fig 4, Table 3). This trend was apparent in the conservative set both when comparing trait-specific eQTL and when comparing eQTL among traits (Fig 4, Table 3). In the permissive set of trait-associated transcripts we observe much more extensive co-localization both within and between traits as well as between male pulse rate and female pulse rate preference (most notably on LG5). Some of the eQTL locations also correspond to or are closely linked to pQTL locations discussed in the previous section (pulse rate: LG 1, LG 3, LG 5, LG 12; carrier frequency: LG 3; pulse rate preference: LG 3 and 5; Fig 3, Fig 4). However, there are also linkage groups that have eQTL for transcripts associated with a trait for which there was no pQTL on the linkage group.

## DISCUSSION

Behavioral barriers to gene flow arise through differentiation in male and female mating communication traits and are a powerful mechanism to promote divergence in the earliest stages of speciation (Coyne and Orr 2004). To better understand this process, we need to examine how the genetic architecture of signaling and preference traits as well as the shape of preference functions and their effect on signal distributions contribute to generating linkage disequilibrium (LD) between co-evolving male and female traits. In this study, we jointly examined the genetic architecture of male signal traits (song) and female preferences in two species of North American field crickets *Gryllus rubens* and *G. texensis*. These species have diverged ~ 0.5 million years ago followed by a long period of bidirectional gene flow that lasted until ~ 18,000 years ago (Blankers et al. 2018b). Preference functions for pulse rate closely track male song distributions, and both male and female traits have diverged conspicuously between the species (Gray and Cade 2000; Blankers et al. 2015b,a).

Our results reveal physical linkage between two co-evolving song traits, pulse rate and carrier frequency, as well as between co-evolving male pulse rate and female pulse rate preference. However, the pQTL of largest effect was never shared between any two traits. We extended our analysis of the genetic architecture into the regulatory pathways that potentially underlie the behavioral traits of interest. We observed tight linkage of eQTL for multiple transcripts associated with the same trait as well as for transcripts associated with different (male and female) traits. This intriguing result suggests linked regulatory variation may contribute to co-evolution of song and preference. Thus, there are multiple dimensions by which physical linkage may contribute to maintaining LD between signals and preferences. However, because physical linkage is incomplete, the striking co-evolution of male and female traits is likely aided by sexual selection resulting from the shape of the pulse rate preference function in relation to the male signal distribution. We hypothesize that these mechanisms jointly facilitate trait-preference co-evolution and the maintenance of a strong pre-zygotic reproductive barrier despite gene flow.

### Integrated song signals

We showed strong, positive genetic covariance between two male song traits, pulse rate and carrier frequency, that are known to be strongly correlated phenotypically (Blankers et al. 2015b, 2017). The strong covariance observed here would allow for a correlated response to selection Although we also report pQTL that are unique to only one trait, the overlapping QTL on LG 1 and LG 3 have relatively high effect sizes (> 10% of difference between species and > 5% of the backcross variance), suggesting that phenotypic effects of linkage may be substantial. This linkage may have resulted in indirect selection on carrier frequency due to strong selection on pulse rate. This process would result in the co-evolutionary patterns observed for carrier frequency and pulse rate across closely related *Gryllus* species, despite no differentiation in female preference for carrier frequency (Blankers et al. 2015a; Hennig et al. 2016). Physical linkage between loci underlying the traits of an integrated sexual signal potentially facilitated signal divergence in multiple dimensions (pulse rate and carrier frequency) even though preference has diverged only in one dimension (pulse rate).

### Integrated features of pulse rate preference

Aspects of female preference are also tightly linked and are likely to co-segregate. It may seem trivial that pQTL scans for pulse rate peak preference and pulse rate preference function (i.e. LD1) are concordant and that focus should be on the discordance instead of the similarity. However, the two measures incorporate different aspects of mate choice. The peak preference score is determined solely by the stimulus eliciting the strongest phonotactic response. This is what in theoretical literature of sexual selection and mate choice behavior is generally considered ‘preference’ (e.g. *sensu* Edward 2015). The linear discriminant function captures multiple aspects of the preference function shape (e.g. peak, width, skew; Fig S1, Table S2). Thereby, this measure is a composite representation of the preference function. The fact that the pQTL scans associated with these distinct measures of preference gave qualitatively similar results, differing only in the magnitude of correlation between genotypes and phenotypes and the presence of a small-effect X-linked QTL, showed that the genetics of peak preference and preference to all tested stimuli are highly integrated. This provides rare empirical evidence for the idea that the genetic underpinnings of difference aspects of mate preference (e.g. peak preference and choosiness, responsiveness) cannot be straightforwardly separated (Kopp et al. 2018).

### Signal-preference co-evolution

Genetic covariance between signal and preference is expected if traits co-evolve within populations and necessary for sexual selection to drive phenotypic divergence (Fisher 1930; Lande 1981; Kirkpatrick 1982; Kirkpatrick and Hall 2004). Empirically, it is unclear whether the dominant mechanism of genetic covariance is physical linkage (proximate loci or pleiotropy) or directional mate preference. Theoretically, both mechanisms would lead to LD between signal and preference and accentuate effects from directional selection (Andersson and Simmons 2006), but LD without physical linkage has been shown to be sensitive to gene flow between partially isolated populations (Servedio and Boughman 2017; Kopp et al. 2018). However, the extent to which physical linkage is required to maintain LD between signals and preferences is also sensitive to the shape of female preferences: e.g. speciation proceeds more readily with open-ended or relative preferences or with strongly divergent unimodal preferences (Kondrashov and Kondrashov 1999, Doebeli 2005).

Here, we show that in *G. rubens* and *G. texensis*, for which detailed demographic analysis have demonstrated divergence in the face of (primary) gene flow, there is some physical linkage between song and preference loci, but also pQTL unique to each trait. Linkage was always between a significant pQTL and a suggestive pQTL or between 2 suggestive pQTL, which are of comparably weak phenotypic effect and associated with more statistical uncertainty. Compared to previous examples in crickets (Shaw and Lesnick 2009), flies (Marcillac et al. 2005), lepidopterans (Kronforst et al. 2006), and fish (Fukamachi et al. 2009) we observe a lesser degree of linkage, showing that divergence in male and female traits does not always require physical linkage (e.g. see Ting et al. 2001; Ritchie et al. 2005; Smadja & Butlin 2009). However, we observe equally strong or stronger levels of genetic covariance between song and preference. We suggest that this is partly explained by the shape of female preference. If the shape of the preference function relative to the population distribution of the signal results in directional selection on the signal, sexual selection can generate strong covariance between traits and preferences without the need for tight physical linkage. In our system pulse rate preferences are non-overlapping, unimodal, and sharply tuned to the male song distribution (Blankers et al. 2015b,a). Comparisons across related *Gryllus* species (Blankers et al. 2015a; Hennig et al. 2016) suggest small differences in the preference result in strong selection on the signal. Depending on the ancestral distributions of the male trait, this may have been enough to drive divergence of signal and preference without strong physical linkage.

We acknowledge there are some caveats to the results discussed here. Statistical (i.e. the Beavis effect) (Beavis 1998) and experimental (selective breeding to optimize phenotypic space in backcross generations) considerations cause our results to be somewhat biased towards loci of large effect. This may either obscure additional linkage among (small-effect) QTL or overestimate the total amount of linkage. Given the phenotypic distances in these closely related species, the sample sizes were not sufficient to detect smaller effect pQTL (< 10%) of which there are likely plenty, e.g.: (Shaw et al. 2007; Blankers et al. 2018a). The fact that only carrier frequency and not pulse rate has X-linked QTL is particularly puzzling, especially in the light of strong signatures of X-linkage for pulse rate in reciprocal interspecific hybrids. This likely reflects the difficulties we had in reconstructing linkage on the X-chromosome because only few markers segregated following expectations for XX-XO mating systems. Additionally, limitations in power to detect pQTL (due to limited phenotypic divergence) make it difficult to distinguish a single pleiotropic locus from multiple loci in close genomic proximity. However, pQTL and eQTL results consistently point towards a mixture of linked and unlinked loci for the co-evolving male and female traits, suggesting that it is unlikely that these caveats have falsely led us to reject completely linked or completely independent segregation of loci.

### eQTL overlap with pQTL

We found strong overlap between pQTL and eQTL, supporting a central role for regulatory variation in behavioral evolution and reproductive isolation (Wray 2007). In some cases, pQTL and eQTL peaks map to proximate or even identical locations. In other cases, we detect more distantly located loci, as well as eQTL on LGs with no pQTL and *vice versa*. One reason pQTL and eQTL might be linked is because both detect a single regulatory variant: many trait-associated SNPs in QTL and genome-wide association studies are regulatory variants rather than protein coding variants (Nicolae et al. 2010) and this is particularly likely for behavioral and sexually dimorphic traits (Wray 2007; Williams and Carroll 2009). For example, small changes in the balance of excitation and inhibition within the neuronal recognition network can rapidly change the phenotype of female preference in crickets and katydids (Hennig et al. 2014). An alternative is that linked pQTL and eQTL represent tightly linked regulatory and coding variants. With the current data, we cannot distinguish between these alternative explanations. The eQTL that did not overlap with pQTL potentially represent true QTL that were not picked up by our pQTL scan, because these loci have only small phenotypic effects but nevertheless sufficiently strong effects on trait-associated gene expression variation to be picked up in the eQTL scan.

### Pleiotropic gene expression and signal-preference co-evolution

The pleiotropic nature of gene expression is well-known as many eQTL detected in transcriptome-wide studies are concentrated in narrow genomic regions (Chesler et al. 2005; Gibson and Weir 2005; Hubner et al. 2005). We detect multiple eQTL for trait-associated transcripts at identical or proximate locations, although most of the co-localization is observed only when the more permissive set of trait-associated transcripts is considered (i.e. all heritable transcripts, including those with lower magnitudes of trait covariation). The most striking co-localization events occur on LG 3, where we detected pQTL and eQTL for transcripts associated with both male song traits and female song preference, and LG 5 where we mapped loci controlling expression of multiple pulse rate and pulse rate preference associated transcripts. We suggest that linkage of regulatory variants reflects an underappreciated genetic mechanism that can affect linkage disequilibrium between signals and preferences. There is limited theory explaining the effects of regulatory variation on the efficacy of sexual selection in the face of gene flow. Existing theory generally indicates that regulatory variation can enhance the effectiveness of assortative mating (Ten Tusscher and Hogeweg 2009) and that linked *cis* regulatory loci can enhance the evolution of sex-biased gene expression (Williams and Carroll 2009) and sexual dimorphism (Connallon and Clark 2010). Our findings provide important empirical insight into the potential for physical linkage, shared regulatory variation, and mate preferences to reciprocally shape LD during divergence with gene flow. We suggest that behavioral, quantitative genetic, and gene expression data be more broadly integrated to understand the effects from sexual selection on diversity across different biogeographic contexts of speciation.

## ACKNOWLEDGEMENTS

We thank Isabelle Waurick for technical support in the lab, D.A. Gray for help with collecting crickets, and Anja Westram and Mingzi Xu for comments on earlier versions of this manuscript. We further acknowledge the support from the r/QTL user group, in particular Karl Broman, and the Biostars and stackoverflow communities. The performed experiments comply with the "Principles of animal care", publication No. 86-23, revised 1985 of the National Institute of Health, and also with the current laws of Germany. The authors declare no conflict of interest. This study is part of the GENART project funded by the Leibniz Association (SAW-2012-MfN-3)

## SUPPLEMENTARY FILES

Table S1. Typical stimulus array for G. rubens female to test pulse rate preference.

Table S2. Correlation coefficients for each of the eight test stimuli in the pulse rate test (Table S1) with their corresponding pulse rates.

Table S3. Genetic map for G. rubens x G. texensis.

Table S4. Transcripts with expression levels correlated to pulse rate.

Table S5. Transcripts with expression levels correlated to carrier frequency.

Table S6. Transcripts with expression levels correlated to pulse rate preference.

Table S7. Dense linkage map.

Figure S1. Preference functions for pulse rate stimuli played back on the trackball system.

Figure S2. LOD score profiles of marker associations with all four traits for single QTL interval mapping.

Figure S3. Heatmaps for between species differential expression.

Figures S4-S6. Heatmaps for LOD scores along the genome for each transcript associated with pulse rate, carrier frequency, and pulse rate preference, respectively.

